# The degree of context un/familiarity impacts the emotional feeling and preaware cardiac-brain activity: a study with emotionally salient naturalistic paradigm using DENS Dataset

**DOI:** 10.1101/2021.08.07.455496

**Authors:** Sudhakar Mishra, U.S. Tiwary

**Author notes:** Corresponding author (S. Mishra); (U.S.Tiwary).

## Abstract

Emotion experiments with naturalistic paradigms are emerging and giving new insights into dynamic brain activity. Context familiarity is considered as an important dimensions of emotion processing by appraisal theorists. However, how the context un/familiarity of the naturalistic stimuli influences the central and autonomic activity is not probed yet [check it]. Hence, we tried to address this issue in this work by breaking it down into three questions. 1) What is the relation between context un/familiarity with the neural correlates of self-assessment affective dimensions viz. valence and arousal; 2) the influence of context un/familiarity in cardiac-brain mutual interaction during emotion processing; 3.) brain network reorganization to accommodate the degree of context familiarity. We found that the less-context familiarity is primarily attributed to negative emotion feeling mediated by lack of predictability of sensory experience. Whereas, with high-context familiarity, both positive and negative emotions are felt. For less-context familiarity, the arousal activity is negatively correlated with EEG power. In addition, the cardiac activity for both high and less context familiarity is modulated before the reported self-awareness of emotional feeling. The correlation of cortical regions with cardiac activity and connectivity patterns reveals that ECG is modulated by salient feature during pre-awareness and correlates with AIC and conceptual hub in high-familiarity. Whereas, for the low familiarity, the cardiac activity is correlated with the exteroceptive sensory regions. In addition, we found that OFC and dmPFC have high connectivity with less-context familiarity, whereas AIC has high connectivity with high-context familiarity. To the best of our knowledge, the context familiarity and its influence on cardiac and brain activity have never been reported with a naturalistic paradigm. Hence, this study significantly contributes to understanding automatic processing of emotions by analyzing the effect of context un/familiarity on affective feelings, the dynamics of cardiac-brain mutual interaction, and the brain’s effective connectivity during pre-awareness.

## 1. Introduction

Mostly, emotion research is done in the lab setting with experimental paradigm lacking ecological validity. Recently, a shift in emotion research using naturalistic paradigm is adapted to probe the cortical dynamics[38, 34, 35]. The stimuli in naturalistic paradigm includes virtual reality immersive experience [25], emotional and neutral movie clips [55, 34], and emotionally salient films [20]. The naturalistic paradigm offers the probe to dynamically track the task-related neural responses, autonomic responses and subjective feedback in a more real, dynamic and natural environment [35]. In emotion research, using a naturalistic paradigm can extend the understanding beyond what is contributed by experiments in the literature with controlled experimental settings (experiments with artificial stimuli lacking ecological validity). In this work, we are probing the effect of affective context un/familiarity in emotion processing in terms of neural activity, autonomic activity and subjective behavioural responses.

The uncertainty in marriam-webster is defined as the lack of knowledge about an outcome or result. Similarly, the state or condition in which something (e.g., the probability of a particular outcome) is not accurately or precisely known is defined as the state of uncertainty in the APA dictionary of psychology. Considering these definitions, if an individual is not able to predict or be less certain about what could be the next sensory experience due to lack of knowledge or context-familiarity, s/he will be in the state of uncertainty. Lack of comprehensive knowledge or context-familiarity promoting difficulty in comprehending multiple possible causes or outcomes is characterized as one possible source of uncertainty [5]. This work explores the uncertainty attributed to less-context familiarity and its influence on central and autonomic dynamics.

Uncertainty and affect are interlinked phenomena that can be mediated by several factors, including information seeking and the subjective perception of lack of information [5]. In literature, many probable theories are proposed to establish this interlinked relationship between uncertainty and affect. For instance, appraisal theorists of emotion processing argued that the degree of context-uncertainty is the fundamental determinant of what specific affect is elicited in a particular situation [37]. The appraisals can be an automatic associations matching with the patterns in the environment. As per these subjective associations, people can have different affective responses to the same situation [37]. Other theories describing uncertainty-affect relationship, including behavioural inhibition system theory (BIS) [18], entropy model of uncertainty [24], and fear of unknown theory [9], have one thing in common that they all view uncertainty as a deficit of knowledge that is inherently aversive and results in negative affective states.

Why the negative affects are prioritized in an uncertain situation? The evolutionary account of the answer proposes that biased prioritization to negative outcomes reduces the cost of missing a potential threat to survival. Hence, the aversive behaviour during uncertainty became the fundamental feature of the mind [8]. For instance, even when a neutral face is paired with the negative gossips, it dominated the activity in the visual cortex and consciousness compared to the pairing with positive and neutral gossips [4]. Similarly, if the perceived stakes of missing the threat are high, it could be advantageous to classify more situations as threats [31]. Even though these evidence sketch an arena associating the negative feelings with the uncertainty, it is unclear what could be mediating this negative feeling and how it emerges out of uncertainty. In this line, a computational model of valence is recently proposed, which states that negative feeling in a less-predictable environment (or uncertain environment) results from fluctuations in confidence in selecting action model. This fluctuation is tracked using affective charge variable (AC). More the mismatch between predicted outcome (generated using simulated action model) and the observed outcome, less confident the agent will feel and consequently will infer negative valence in the particular situation [23]. However, this computational model is still lacking the empirical support, which we are providing in this work.

The brain controls the rate, rhythm, and power of heart-beats via both sympathetic (‘fight and flight’) and parasympathetic (‘rest and digest’) axes of the autonomic nervous system [54]. In addition, cardiac functions can be profoundly altered by the reflex activation of cardiac autonomic nerves in response to central autonomic commands associated with stress, physical activity, and arousal[52]. For instance, the increased attention to suppressing the uncertainty down-regulated the tonic HR and high-frequency HRV. This cardiac deceleration was sustained in binocular-rivalry-replay task and modulated by ambiguity, novelty and switch rate [11]. On the other hand, the beat-by-beat operation of the heart is also tuned to engage the emotional and cognitive resources [48]. Additionally, the afferent cardiac signals also facilitates brain activity and modulate the sensory attention to perceive the low-spatial frequency features of fearful faces [6]. For instance, the cardiac activity, mediated through the amygdala, facilitates fine-grained discrimination at the sensory cortices [56] and promotes active sampling to resolve the uncertainty [3]. Conforming to this two-way interaction, a computational model based on active interoceptive inference (AII) theory postulates the influence of cardiac arousal activity on exteroceptive sensory processing as well as modulation of the cardiac activity when the simulated agent is presented with the arousing stimuli as compared to the relaxing stimuli [3]. The empirical support to this model with ecologically valid experiments is still lacking. Primarily, in the literature, the cardiac-brain mutual interaction is probed with the experiment paradigm lacking in ecological validity [11, 6]. Hence, how the cardiac and brain activity mutually interact with varying degrees of context familiarity during the pre-awareness period is not yet probed. In addition, the underlying automatic mutual dynamics to resolve uncertainty are also not apparent during this time [5]. Aiming at these gaps, this work extends the understanding of cardiac-brain interaction dynamics by addressing how the cardiac activity interacts with the neural activity to play a constitutive role in emotion processing amid varying degrees of context un/familiarity in an ecologically valid naturalistic setting.

To this aim, we are asking the following questions in this work. 1. How the context-un/familiarity is dynamically correlated with affective psychological dimensions? 2. How does the degree of context un/familiarity affect the cardiac-brain mutual interactions? 3. How the change in neural activity is associated with the context un/familiarity?

To address the questions described above, we have designed an EEG experiment with an ecologically valid experimental setting (see section-2.1). The raw signal is preprocessed and analyzed as follows. 1) behavioural analysis on subjective ratings; 2) correlation analysis between subjective ratings and power of different brain networks; 3) frequency analysis of cardiac activity; 4) correlation analysis between ECG power and brain networks’ power; 5) classification of high-context familiarity from less-context familiarity using machine learning; 6) phase synchronization analysis to find out functional connectivity and to find out cortical regions with high-centrality value; and 7) effective connectivity calculation using granger causality between nodes with high centrality value.

We found that uncertainty is associated with negative affect and this affective feeling is mediated by low confidence in prediction about the sensory experience. Additionally, uncertainty is associated with high arousal and EEG desynchronization promoting information seeking and adaptation to the new information. Analysis of cardiac-brain activity reveals that cardiac activity is more correlated with exteroceptive sensory regions and facilitates information-seeking during less-context familiarity. Moreover, uncertainty engages the cortical regions (OFC & dmPFC), which tracks the changes and adapt to the dynamically changing situation. We present the results and discuss these findings.

## 2. Methodology

### 2.1. Experiment

We have designed an EEG experiment along with the ECG recording. We used multimedia stimulation (selected after a carefully designed stimulus validation process as described in [34]) in our experiment. A total of 40 participants participated in this experiment with mean age (23.35 ± 1.34) and three female participants. All were enrolled in the institute in the bachelor and master courses (for detail see [34]).

The experiment paradigm is shown in the fig-1a. Participants are well informed and trained for the emotion task. While watching the naturalistic stimuli (whole paradigm is detailed elsewhere [34]), participants did the mouse click to mark the emotional feelings. They can click any number of times. After each stimulus, participants are presented with feedback scales: valence, arousal, dominance, liking, familiarity, and relevance. Participants are instructed to rate the familiarity scale as follows. “While rating the familiarity scale, recall how well and frequently you were able to predict what could be the next event or episode or scene in the movie clip. According to this self-judgement, you have to rate the familiarity scale from 1 (less-familiarity) to 5 (high-familiarity)”. Additionally, participants labelled the mouse click with the emotion categories. To help in recalling and labelling emotional categories, participants are shown the frames extracted around the mouse clicks. With this simple trick, we were able to localize emotional feelings of the participants.

**Figure 1:**
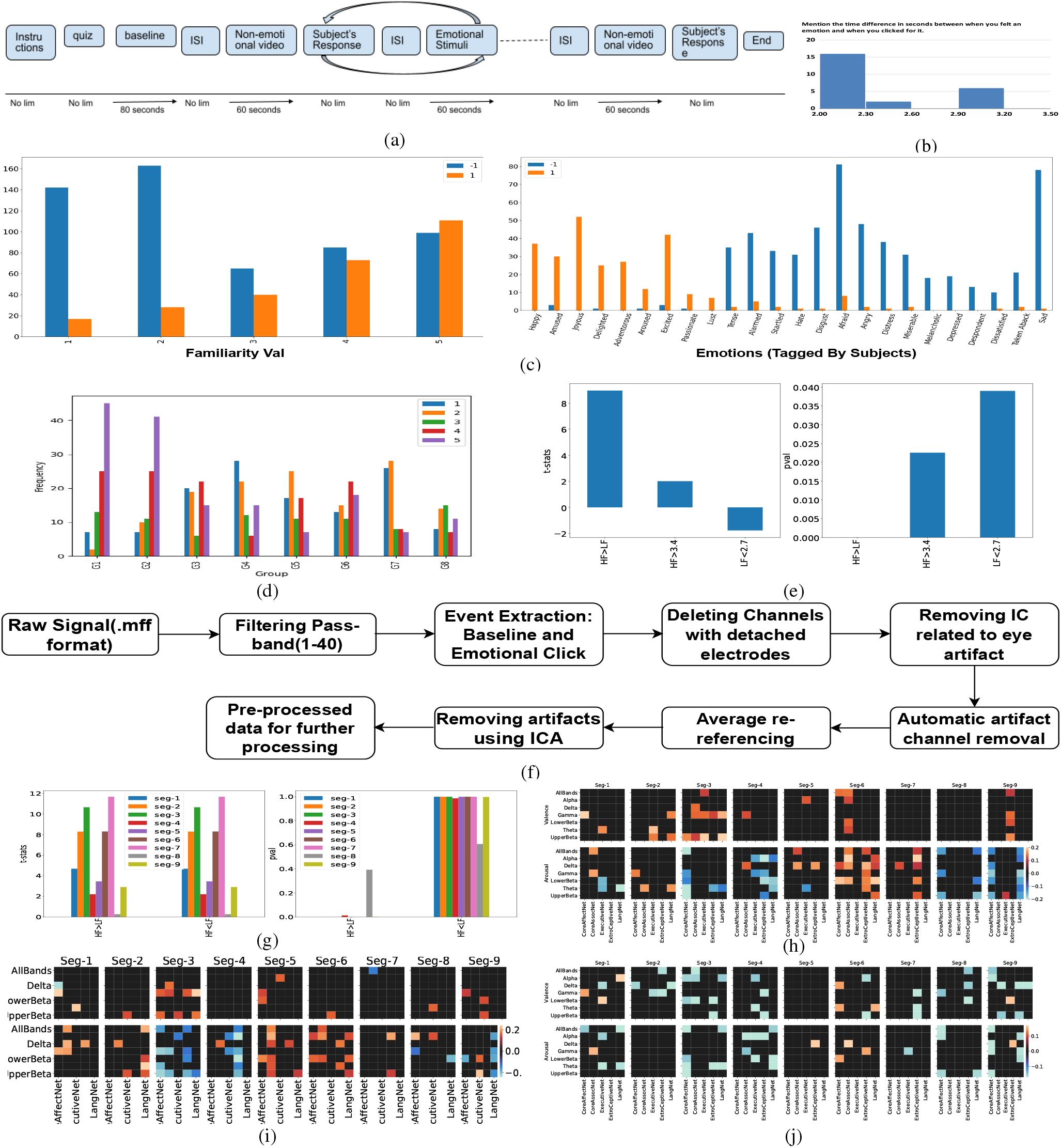
(a) Experiment paradigm (b) Time duration before click when participants consciously responded to their feeling of any emotion. (c) Left: histogram of familiarity ratings associated with negative and right: positive feelings, histogram of 24 emotions (as reported by subjects) with their positive and negative associations, (d) The distribution of familiarity ratings in the scale 1(low familiarity) to 5(high familiarity) for all the emotion groups. Groupings are shown in the supplementary fig-1. (e) Paired ttest with the alternative hypothesis that familiarity ratings for HF group are greater than LF group. Moreover, results of one sample ttest with the population mean 3.4 and 2.7 for HF and LF groups, respectively are shown. The high-familiarity ratings are greater than 3.4(*p <* 0.025) and less-familiarity ratings are less than 2.7(*p <* 0.04) (f) Preprocessing pipeline. (g) Segment-wise paired ttest with the alternative hypothesis that average power of EEG channels for HF is greater than LF (left) and LF is greater than HF (right). The power for LF group is significantly lower than HF group except seg-8. (h-j) Represents the significant correlations calculated (with *p <* 0.05) between the valence and arousal subjective ratings and average normalized power for different brain networks [eeg channels are source localized to brain regions; for all the emotion groups(h), high-familiar group(i) and less-familiar group(j) are considered.]

### 2.2. Pre-Processing

The preprocessing steps are shown in fig-1f. The raw signal is imported, which is already referenced to Cz(reference electrode, EGI default). However, we performed average rereferencing during the preprocessing. The raw EEG signal is filtered with a fifth-order Butterworth bandpass filter with the low-cutoff 1 Hz and high-cutoff 40 Hz. From the filtered signal, the event segments corresponding to baseline state, pre-stimulus, and the click are extracted with time-duration 10s to 70s, −3s to 0s, and −6s to 1s, respectively. The extracted duration for the click event is from 6s before the click event to 1s after the click.

Next, the concatenated signals were manually checked. The channels and samples with very high amplitude, probably caused by electrodes detachment, are removed manually (The manual rejection sheet is online). Then, we applied the ICA to remove the eye blink artefact so that high amplitudes of eye artefacts should not misguide our next step, which is automatic channel rejection. Then, the second ICA is applied for any other remaining artefact in the signal (including eye artefact, muscle artefact, heart artefact, line artefact, and channel artefact). We kept independent components with the probability of brain activity more than 0.3 (adding the probability factors for all the artefacts, including brain signal, equal to 1). After completing the preprocessing steps, the individual events corresponding to baseline state and emotion clicks are stored in .mat files for further data analysis.

For ECG, the channel data is extracted and bandpass filtered with low-cutoff 0.6Hz and high-cutoff 40Hz (all other filter parameters are the same as with the EEG processing). Only segments corresponding to the time duration for the resting state event and click event (duration mentioned in the above paragraphs) were considered from the continuous signal. The ECG signals are shown in supplementary fig-2.

### 2.3. Source Localization

The preprocessed and extracted EEG segments corresponding to each click event are used for our source localization procedure. In this study we have usedsLORETA) method [41]. It is a distributed inverse imaging method. The current density estimate is based on the minimum norm (*l*_2_ − *norm*) solution, and localization inference is based on standardized values of the current density estimates. sLORETA is capable of exact (zero-error) localization. The objective function to be minimized to get zero error localization is

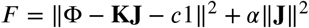

where *α* ≥ 0 is a regularization parameter. This functional is to be minimized with respect to **J** and c, for given **K**, Φ and *α*. The explicit solution to this minimization problem is

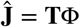

where:

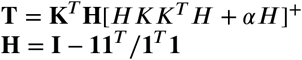

with 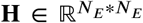 denoting the centering matrix; 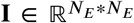 the identity matrix; and 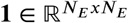 is a vector of ones.

### 2.4. Seed-Coordinates from meta-analysis and Emotion related networks

All the processing is done in MNI voxel coordinates. MNI152 template is used for the source localization (5×5×5mm voxel size), and anatomical labelling is done as per the sheet provided by the sLoreta software [41]. This sheet had 6239 voxels with size (5×5×5mm). We selected voxels corresponding to those regions which were reported in the meta-analysis study for emotion [29]. In the meta-analysis [29], regions that were activated across emotion studies are grouped into five networks including, core-Affective, core-Associative, language, executive control, exteroceptive sensory. Regions along with original coordinates are attached online. Since the reported coordinates in the study [29] are with the voxel size (1×1×1mm), and we have voxels with size (5×5×5mm), we calculated all the voxels within 5mm Euclidean distance from the coordinate originally reported in the meta-analysis study. This gave us a new set of voxels (see the attached sheet recalculateVoxels.xls). Finally, we got 216 voxels.

### 2.5. Correlation of Power of brain networks with the Valence and Arousal ratings

The 7 seconds signal is divided into eight segments (without zero padding) with 250 samples per segment and an overlap of 75 samples between the segments. For each frequency band and segment, average power for brain networks is calculated using the welch power method with Hanning window (to deal with edge artefacts due to segmentation of the signal).

We calculated the Pearson’s-r correlation between the average normalized power (stimuli power divided by baseline-state power) and the self-assessment ratings (viz. valence and arousal) with significance value of 0.001 for each network, frequency band and segment. Results are shown for all emotion groups together (fig-1h), for high-familiar group (fig-1i) and for less-familiar group (fig-1j).

### 2.6. Phase locking value Connectivity and Network Profile

For each frequency band and each seg, the phase-locking value (PLV) is calculated among 216 voxels (constituting five networks). The real signal is transformed into an analytic signal using the Hilbert transform. Phase and amplitude envelope is calculated from the analytic signal. Functional connectivity among all the voxels using phase synchronization is calculated as follows.

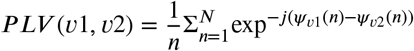

The significance of the functional coupling is calculated using the Wilcoxon-sign-rank test with *p* − *value <* 0.001. We calculated these significant connections individually for each emotion group, segment and frequency band.

### 2.7. Centrality Calculation

The centrality calculation (for finding hubs) is done for each frequency band and for each segment. Hub is one of the global metric of complex networks which can be found out by calculating centrality value for each node. We calculated betweenness centrality value for each node(voxel) using the following equation:

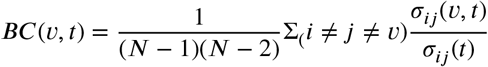

where *σ*_*ij*_(*t*) is the total number of shortest paths from node i to node j at time t and *σ*_*ij*_(*v, t*) is the number of those paths passing through v at time t.

Betweenness centrality quantifies the extent to which a node participates in the shortest paths throughout the network. Nodes with high betweenness centrality are thought to be particularly influential across different efficient pathways of the network as a whole rather than just local direct connection. To reduce the chances of considering false centrality values, we created 100 random graphs for 54 pairs (6 frequency bands x 9 segments). These random graphs are created with the same number of voxels simultaneously, preserving the degree distribution. The mean threshold (across all the conditions) for each voxel is calculated as follows:

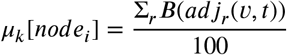

where k=1:54 (54 conditions) and r=1:100 (100 random graphs ∀ condition indexed as k). *adj*_*r*_ is the adjacency matrix for random graphs. B symbolized betweenness centrality calculation. *µ*_*k*_[*node*_*i*_] is the vector containing the mean centrality value for each node calculated over random graphs for each condition.

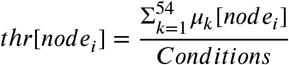

*thr*[*node*_*i*_] global threshold for each node. This global threshold is used to select nodes with a centrality higher than this global threshold. The calculated hubs with their centrality value are shown (supplementary fig-4).

### 2.8. Granger Causality

G-causality is based on predictability and precedence. A variable X is said to G-cause a variable Y if the past of X contains information that helps predict the future of Y over and above the information already in the past of Y. In this article, the Granger causality is calculated using Multivariate Granger Causality (MVGC) MATLAB toolbox. The G-Causality is formalized using vector auto-regressive modelling. The calculated G-causal connections were tested for significance against the null hypothesis of zero causality using the F-test. Further, the obtained p-values from F-test are corrected for false discovery rate with the significance level *α* = 0.01.

### 2.9. ECG Analysis

ECG is recorded simultaneously with EEG. We filtered the ECG signal with the pass-band from 0.5Hz to 40Hz. The obtained ECG signal is shown in (supplementary figure fig-2). The obtained ECG signal is divided into two parts. The first part ranges from 0 to 600 samples 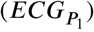, whereas the second part ranges from 600 to 1200 samples 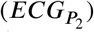. We transformed the time domain signal into the frequency domain using Fourier transform and then calculated the total power for frequency range 1-15Hz (low-frequency range) and 15-40Hz (mid-frequency range). We calculated the significant change in power between the first and second parts for low and mid-frequency using the right tail student t-test (with the 95% confidence interval; tab-2b). For high and less context familiarity, we performed the right tail student t-test with the 95% confidence interval on sub-sample means (30 samples are randomly taken from the total set of samples for high-context familiarity and less-context familiarity; tab-2c; mean distribution is shown in supplementary fig-3).

Next, we calculated segment-wise Pearson’s correlation between power of different brain networks with ECG power (calculated for low and mid-frequency range). We checked the significance of correlation with two significance levels: 0.01 (denoted with #) and 0.05 (denoted with). See fig-2a, 3a, 3b for all emotion groups, for emotion groups with high-context familiarity and less-context familiarity, respectively.

**Figure 2:**
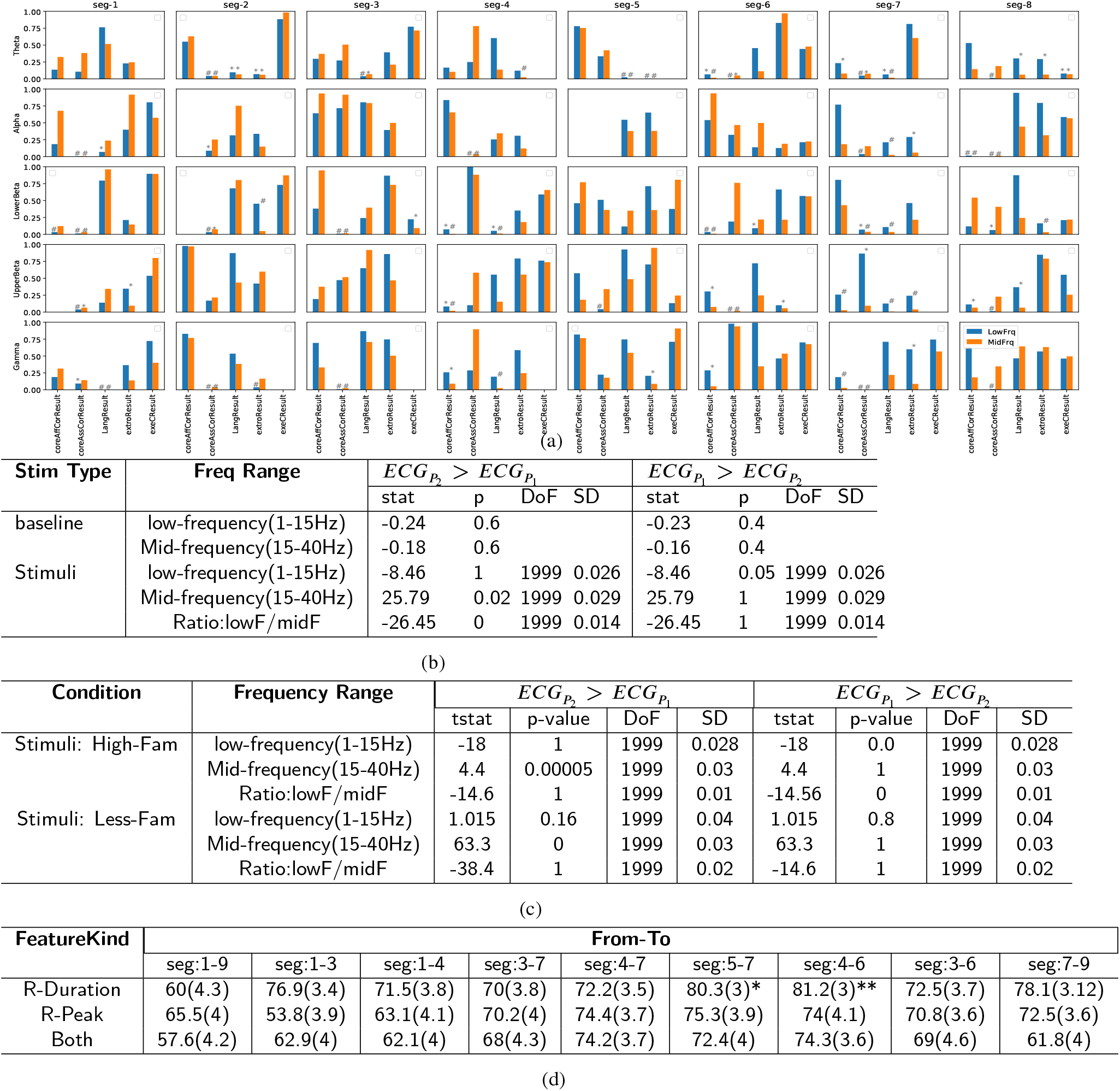
(a) Pearson’s correlation test for power of different brain network with ECG power (all emotion groups are considered). # and denotes significance level 0.01 and 0.05, respectively. (b) Wilcoxon signrank test with 5% signiicance level and alternative hypothesis that 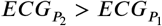 (third column) and 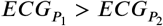 (fourth column) (c) Same test as (b) for emotion groups with high-context familiarity and less-context familiarity. Mean distribution for HF and LF is shown in supplementary fig-3. (d) The table is representing the accuracy for the k-nearest neighbour classification between the high-familiar and low-familiar classes. R-duration and R-peak are the duration of the R wave and the amplitude of the R wave.

**Figure 3:**
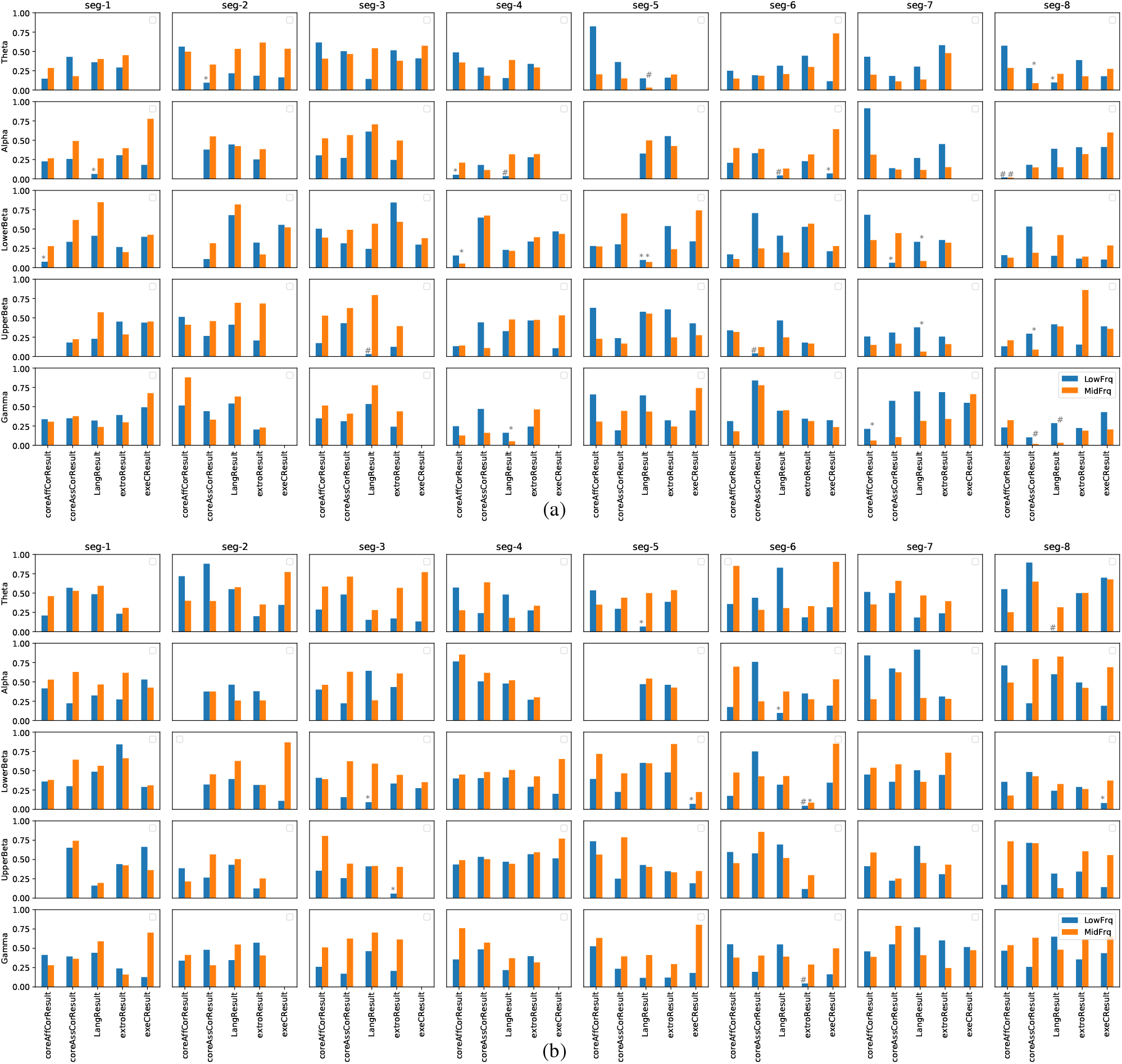
Pearson’s correlation test for power of different brain network with ECG power. # and denotes significance level 0.01 and 0.05, respectively. (a) high-context familiarity groups and (b) less-context familiarity groups

### 2.10. Equations

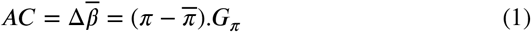

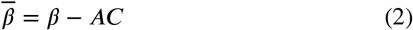

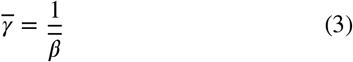

where AC is the affective charge which compares the prior and posterior policies and tracks fluctuations in expected precision, hence, confidence in expected action model *G*_*π*_. *π* and 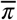 are prior and posterior policies, respectively, which are formulated in terms of baseline prior(*E*_*π*_), expected free energy(*Gπ*) and perceptual evidence(*F*_*π*_) as follows:

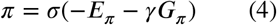

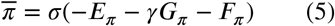

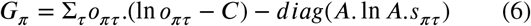

Where, *o*_*πτ*_.(ln *o*_*πτ*_ − *C*) is the expected phenotype risk and *diag*(*A*. ln *A*.*s* _*τr*_) is expected perceptual ambiguity. For the treatment of full formulation readers is suggested to read[23].

## 3. Results

### 3.1. High and less context familiarity groups

The grouping of emotions is described in [35] and supplementary fig-1. The subject has rated their familiarity in the familiarity scale ranging from 1(low-familiarity:LF) to 5(high-familiarity:HF) for each emotional stimuli. These subjective scores on familiarity are grouped and presented in the form of a histogram (fig-1d). Emotion groups-1,2,3 & 6 are inclined towards the HF group in comparison to emotion groups-4,5,7 & 8, which are inclined more towards the LF group. To test this observation, a two-sample t-test is performed. Ratings for the HF group are higher than LF groups with significance (t=8.97 & *p* = 1.7*e*^−18^) (fig-1e). The follow-up question is what are the significant thresholds for HF and LF groups. A one-sample t-test with the population mean 3.4 and 2.7, respectively, for HF and LF groups is performed. HF groups are greater than the mean 3.4 with the significance (t=2;*p <* 0.025;fig-1e). Whereas, LF groups are lesser than the mean 2.7 with the significance (t=-2;p<0.04;fig-1e).

### 3.2. Less-context familiarity leads to negative feelings and asynchronous brain activity

For HF groups(fig-1i), the valence values are significantly correlated with the brain networks in the upper beta and gamma bands in seg-3 followed by very less/no significant correlations in the coming segments. On the other hand, the negative correlation of arousal ratings is followed by a positive correlation with the brain networks. Our interpretation is that subconscious detection of salient features during seg-3 and seg-4 caused increased attention [14] to stimuli and desynchronized brain activity [57, 50] followed by high correlation of arousal (seg-5 & seg-6) which is related with synchronized oscillatory power. On the contrary, if the context is unfamiliar (or less-familiar), the correlation between valence ratings and EEG power is negatively correlated at seg-3, and seg-4 with the regions in core-Affect, core-Associative and exteroceptive networks, followed by relatively positive correlation with core Affect and negative correlation with the exteroceptive sensory network. So, compared to high-context familiarity, the valence correlated neural activity fluctuates throughout the pre-aware period. If the context is less familiar, the subject is trying to figure out the value of the stimuli and is inclined towards negative emotions. For both HF and LF groups, during the seg-3 and seg-4, the arousal activity is negatively correlated. This effect is mediated by change in perception due to detection of salient feature in both conditions. In addition, low power and correlation during seg-5 & seg-6 in LF group, hints the uncertainty in perception, hence, desynchronization. This result is additionally confirmed with the analysis of segment wise power comparison for HF and LF groups. The segmentwise power of eeg electrodes is significantly greater for HF group than LF group for all the segments [except seg-8] (fig-1g). One of the reasons for low power is asynchronous brain activity due to less-context familiarity [59]. The asynchronous irregularity is associated with high arousal.

### 3.3. Affect modulates cardiac activity

The next question was, how the cardiac-brain mutual interaction takes place during the pre-awareness? To this aim, we have calculated the correlation of ECG power with the EEG power for different brain networks, segments and frequency bands. We found a significant correlation between the core-Affective network and ECG from seg-4 to seg-7. During the seg-4, ECG power is correlated with the core Affect (mainly Ofc and AIC fig-2a & 4a; lower beta and gamma bands) and language (mainly ATL and VLPFC fig-2a & 4a; alpha, lower beta and gamma bands) related brain regions is observed. This correlation continued till seg-7 in theta, lower beta, upper beta and gamma bands. Mostly, the modulation is observed in the mid-frequency range for both the core-Affective and language networks (fig-2a), which is in sync with our subsequent correlation analysis.

**Figure 4:**
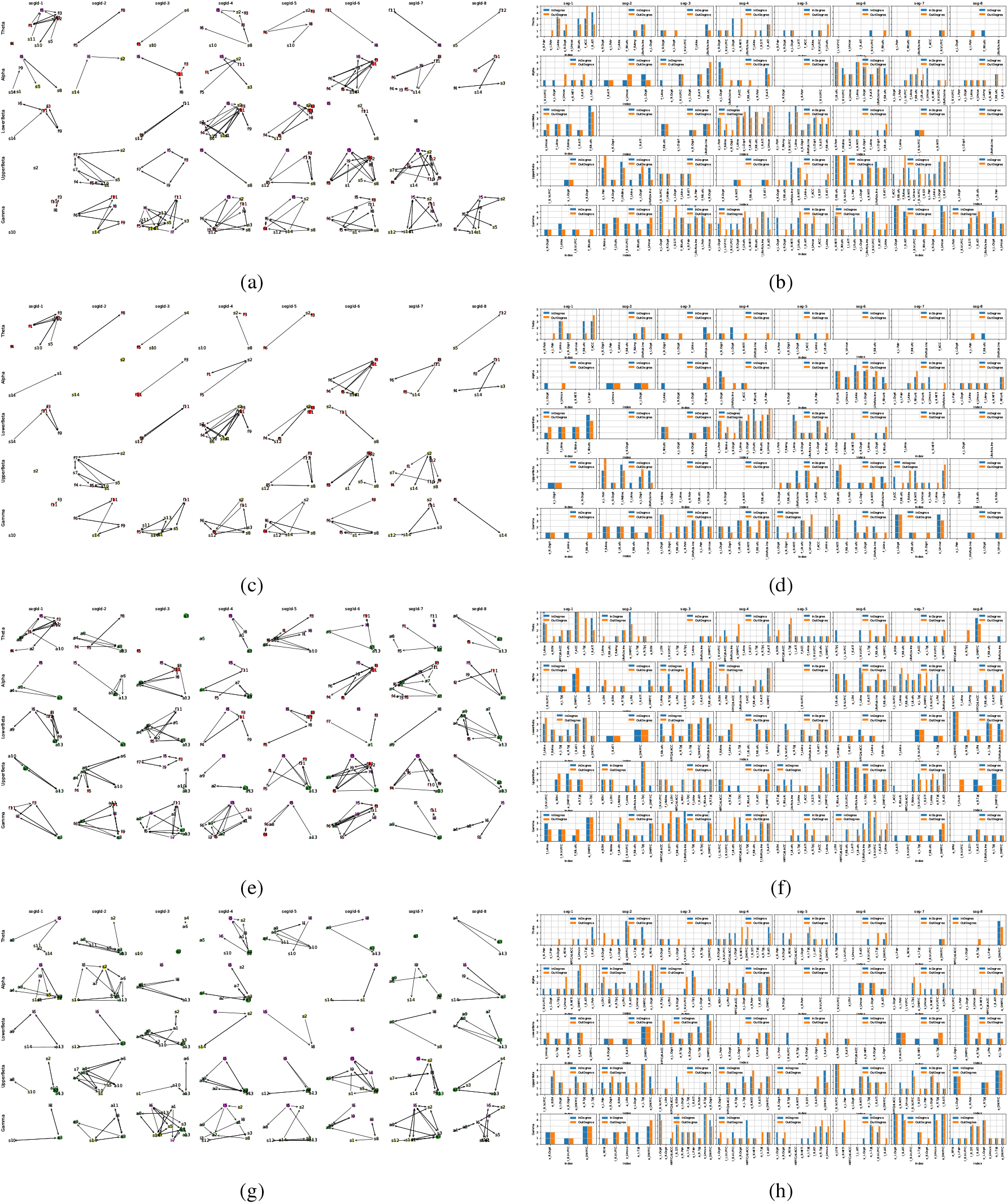
All the graphs are created for the representative (median) connections across the emotions (left) and degree histogram for groups (right). (a-b) For language, core Affect and exteroceptive sensory related regions. (c,d) For core Affect and exteroceptive sensory related regions. (e-f) For language, core Association andcore Affect networks. (g-h) For language, core Association and exteroceptive sensory region. (a,c,e,g) The granger connectivity graph, (b,d,f,h) the calculated indegree and outdegree bar plots. For encoding of regions see supplementary table-1

We further tested the modulation in ECG activity during seg-4 to seg-7 (i.e. 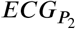). During the baseline state, there were neither significant differences for the low-frequency range nor for the mid-frequency range (tab-2b) between 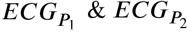 (see section-2.9). On the contrary, for the stimuli condition, we observed a significant difference (*p* − *value* ~ 0.02) in the mid-frequency range for the comparison condition 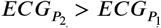. Hence, during the seg-4 to seg-7, the autonomic physiological activity is modulated.

To test the ECG modulation for HF and LF groups, we performed the t-test for the distribution of subsample means for both HF (left in supplementary fig-3) and LF groups (right in supplementary fig-3) with the significance value (p<0.05) (tab-2c). We found a significant increase in power for the mid-frequency range for 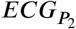 in comparison to 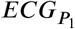. On the contrary, low-frequency power is decreased for 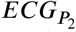 compared to 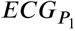. On the other hand, the same pattern is observed for the less-familiar group for the mid-frequency power, whereas no significant change is observed for the low-frequency power. This suggests that during the processing of high-familiar emotion stimuli, the low-frequency power of ECG decreases which is not the case with the less-familiar emotional context.

Furthermore, we analyzed the impact of context un/familiarity on correlation of ECG power with the power of brain networks (fig-3a & 3b). For the HF group, the ECG power is correlated with language, core Affect and association networks (fig-3a). On the other hand, the correlation with exteroceptive sensory regions is observed for the less familiar group, which may be due to foraging for the information to resolve the uncertainty. In addition, the connectivity of the executive control network is increased for LF than HF, which is reported to enhance information gathering and resolve reward uncertainty[17].

We then asked whether ECG can predict the context. Table-2d depicts the accuracy for the k-nearest neighbour (k-NN) classification between high-familiar and less-familiar classes for different combinations of segments. The maximum accuracy is observed for R-width feature during seg-4 to seg-6 followed by seg-5 to seg-7. The R-width is related with the frequency (see supplementary fig-2). For low R-width, high frequencies show more amplitude than the low frequencies and vice-versa. Hence, in sync with the previous analysis the context familiarity information is encoded in the ECG frequency. Notice here that maximum information for classifying HF from LF group is observed between seg-4 to seg-7 which is in sync with the power-correlation analysis (fig-1i & 1j), ECG power-spectrum analysis (tab-2c) and power-correlation analysis between ECG and cortical brain networks (fig-2a, 3a & 3b). Moreover, during this period, the subject is unaware of this autonomic activity (as per the subjects report in tab-1b).

### 3.4. High and low context familiarity engages different brain regions

In the effective connectivity graphs (fig-4), we observed that mid-insula and anterior-insula are receiving inputs from higher order sensory regions such as LOcptl and LPstr in the upper beta band in seg-2 followed by bidirectional activity of AIC with conceptual network nodes in seg-3 (RATL& RVLPFC) (fig-2a, 4a & 4d). During the seg-3, in the gamma band, the STL has directed connectivity to higher order sensory cortices (AIC, LOcpt, and LPstr;fig-4a, 4b) and to the region reported in mental-self activity(DMPFC;fig-4e, 4f). During the seg-3 in the alpha and lower beta band the connectivity of core Affect nodes(LAIC& Rlt.OFC) with mental-self (DMPFC) and self-other distinction(TPJ) related regions is increased. One interpretation is that the subject is trying to infer the relevance of detected salient feature with self.

During the seg-4, the connectivity of RATL with higherorder sensory (LOcptl), interoceptive insula and reward/punishment related node(Rlt.ofc) is increased (fig-4). Overall from seg-4 to seg-7 the unidirectional and/or bidirectional effective connectivity among regions in networks including, language, core Affect, and core Association, is increased (fig-4e&4g). Importantly, effective connectivity between language and core Affect related regions is increased from seg-3 to seg-7; ls & lf Figures reference). It is in sync with our ECG results (fig-2a), where the correlation of ECG power with language and core Affect regions is increased. Hence, a coordinated activity between language and core Affect regions is influencing the autonomic physiological activity.

Next, we probed how the effective connectivity for HF and LF groups is different. For both groups, we calculated the median of effective connectivity values (median network). Following this, the calculation of in-degree and out-degree for the median network is done. The difference between in-degree and out-degree for high and less-context familiarity is calculated (as shown in fig-6). We have observed increased connectivity of OFC for the less-familiar stimuli in the alpha band. Mainly, it is connected with R.ATL and L.AIC (fig-**??**). Increased oscillatory activity in OFC is reported in depressed and negative mood state [45], reward uncertainty, encoding reward-biased and risk-modulated subjective value dynamically[27], and regulating the uncertainty[1]. Whereas, increased insular connectivity, mainly in the lower beta and gamma bands, is observed for the high-context familiarity. AIC performs multimodal bottom-up integration and projects top-down information to the primary interoceptive sensory cortex. AIC is in the path of descending interoceptive information, which provides a point of reference for autonomic reflexes[19]. The dmPFC has relatively more connections with other brain nodes for LF group than for the HF group. The activity in dmPFC is reported in resolving valence conflict[28]. For both HF & LF, activity in the right Anterior Temporal Lobe (rATL) is observed. Anterior temporal lobes are generally involved in social cognition, theory of mind processing, social conceptual knowledge, semantic integration, and memory systems that are linked to reward and valence[49, 22].

## 4. Discussion

### 4.1 Degree of confidence in expectation impacts positive and negative feelings as well as intensity

Biologically plausible forms of cognition and affect inevitably involve policy selection. Stimuli with high familiarity will have high certainty in policy selection and high confidence in the en-activated action model. On the other hand, less familiarity induces perceptual and affective ambiguity. Amid perceptual ambiguity, information seeking and detection of a new significant or salient event updates the posterior beliefs about self-state (i.e. expected action model) which in turn causes the fluctuation in affective charge (1) [23].

The valence is the higher-level state inferred from these fluctuations in AC. AC dynamically tracks the expected precision of the action model (1) given the policies ((4) & (5)) [23]. In the valence row (fig-1i), the correlated fluctuation in EEG power during seg-3 is observed for high-context familiarity, whereas for the stimuli with less-context familiarity (fig-1j), the fluctuations in correlation are more often appearing and each fluctuation corresponds to the new policy selection following the detection of new salient information in order to reduce the uncertainty. These continuous fluctuations also signify the lower confidence in expectation, which is inferred as negative valence[5, 23]. Whereas, in case of high familiarity, the agent is certain about the selected policy and action model hence, very low or no policy mismatch leads to no fluctuation in the affective charge ((1)) and high expected precision ((3)) on the action model(valence row in fig-1i).

During seg-3 & seg-4 for HF (fig-1i) as well as LF (fig-1j), as soon as the salient external information (not matching with the ongoing prediction) is detected, the ongoing prediction about sensory experience is updated and projected on interoceptive and exteroceptive sensory regions (fig-1i). This makes the sensory network activity asynchronous (low EEG power) and sensitive to the afferent input during seg-3 and seg-4 (arousal row in HF:fig-1i and 1j) [53, 59]. On the other hand, for HF, during the seg-5 & seg-6, the matching of sensory expectation with the observation (arousal row in fig-1i) leads to synchronous regular state representing the collective endogenous arousal pattern and getting unresponsive to the stimulus [59]. On the contrary, for less familiar context, during the seg-6, the correlation is low but positive compared to seg-3 and seg-4 (fig-1j). This relatively high correlation (i.e. high EEG power in seg-6, arousal row, fig-1j) is indicating reduced mismatch between expected and observed sensory sampling compared to seg-3 and seg-4 (fig-1j) but still less synchronized state compared to the high-context familiarity during seg-5 & seg-6 (arousal row in fig-1i).

The computational model of valence [23] defines the valence as the inferred state from fluctuations in affective charge in the predictive coding framework. The fluctuations in AC due to prediction uncertainty are reflected in the correlation of valence with the power of electrophysiological recordings. Moreover, in this analysis, we have reported that uncertainty drives a state of high arousal and asynchronous irregularity (AI) [53], which, in turn, is associated with low EEG power. This association of uncertainty with AI is also justified with the predictive coding point of view in a way that AI increases sensory awareness, which can be used to adapt new salient information from the environment to resolve the uncertainty.

### 4.2. Cardiac-brain interaction

For high-context familiarity, we observed that 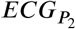 shows higher accuracy in classifying stimuli in HF and LF groups. These results hint that the pre-aware autonomic activity contains the information about context un/familiarity. These results indicate that the spontaneous neural activity (processing affective context) and cardiac activity mutually influence each other even before the subject’s awareness (selfreported awareness around seg-6 & seg-7; tab-1b). These results are supported and further detailed in our power correlation analysis of ECG with the brain networks. In this analysis we found that the higher-order core Affect and language related regions are correlated with the cardiac activity for the high-context familiarity during seg-4. On the contrary, no such activity is observed during seg-4 & seg-5 for less-context familiarity. Instead, for less-context familiarity in seg-6, the cardiac activity is correlated with the exteroceptive regions in theta and lower beta bands which could be related to active sensory sampling facilitated by the cardiac activity [56].

We have found an increased power-correlation of ECG with power of core Affect(AIC & ofc) and language(ATL) related regions (fig-2a, 4). It provides evidence for the association of body physiology with the conceptual hubs and VMA regions. We believe that during high-context familiarity, the already learnt emotion concept is reactivated and the associated representations, including visceromotor representation is simulated to re-experience the feeling of an emotion which is felt in the past with a similar context or event [35]. The learning of emotions with the autonomic coregulation of visceromotor activity starts early in childhood during mother-infant interaction. For instance, in the developmental psychology literature, the autonomic socioemotional reflex (ASR) theory posits that the mother-infant mutual stimulation helps in learning the autonomic conditional reflexes related to positive and negative emotions and corresponding visceral/autonomic co-regulation [30]. ASR describes the approach and avoidant behaviours between mother and infant as ‘autonomic emotional connections’ and ‘autonomic emotional dysconnections’, respectively. Physiologically, these behaviours co-regulate physiological activity such as cardiac reflexes. This autonomic learning helps regulate attention and stress in as early as 12-months old [58]. In addition, during the childhood calming sessions restore a positive ASR and lower baseline heart rate (HR) over time which promotes adaptive socio-emotional growth and development [30]. This adaptive learning learnt through interaction with mother in childhood and interaction with complex social patterns later in life helps to regulate the adapted homeostasis by making predictions inferred from the learnt experiences & concepts in the past. Due to life-long learning of the association between body physiology and brain activity, access to the learnt emotion concept happens automatically before the individual becomes aware of physiological changes. This gives an advantage in reducing metabolic, and allostasis load [26].

In addition to the support from psychology, our results for cardiac-brain mutual interaction is also underpinned by the computational model of active interoceptive inference (AII) [3]. Recently, [3] proposed the first computational framework of active interoceptive inference (AII) in order to probe ‘how ascending cardiac signals entertain exteroceptive sensory perception and confidence?’. Three conditions were simulated for the healthy and lesioned (deafferentation) agents.

In simulation-1 of the AII computational model, the cardiac activity is modulated for the healthy agent as soon as the arousal stimuli are encountered, and this modulation is followed for some duration. A similar modulation we have observed in our ECG modulation analysis where *ECG*_*P*_2 has shown high power for mid-frequency range in comparison to 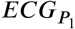 (fig-2b). Additionally, for the high-familiar groups, there is significant interaction of core Affect regions with cardiac activity. However, for less-familiar groups, the cardiac activity is mainly correlated with the exteroceptive net-work (seg-3 & seg-6 in fig-3b). This can be interpreted, in sync with the simulation-2 [3], that cardiac activity had a direct effect on exteroception. Whereas exteroceptive activity had an indirect effect (via policy) on interoceptive activity. For the stimuli with less-context familiarity, cardiac activity mutually interacts with the exteroceptive activity with less or no mediation by interoceptive regions (VMA regions). Hence, no correlation between core Affect regions and cardiac activity is observed. Moreover, we believe that the correlated cardiac activity with exteroceptive sensory regions could be related to active sensory sampling synchronized with cardiac activity. By this active sensory sampling, participants are trying to achieve the fundamental evolutionary imperative, which is “resolving uncertainty” [51].

Our results provide empirical evidence for the computational model of AII in ecologically valid setting with the naturalistic stimuli. In addition, we provide evidence for autonomic co-regulation of cardiac activity with VMA and concept related regions during high context familiarity. On the contrary, the afferent cardiac signals influence the exteroceptive sensory activity to perform active sampling while processing stimuli with less-context familiarity.

### 4.3. Differential processing of higher-order regions for high and less context familiarity

In fig-6 & 4), we observed that higher-order brain regions are contributing differentially. For instance, OFC shows increased connectivity for less-familiar contexts in the alpha band. Whereas, increased insular connectivity, mostly in lower beta and gamma bands, is observed for high-familiar context. In addition, the TPJ connectivity is observed in the theta, alpha and gamma bands for the high-familiar context. Whereas, for the less-familiar context, dmPFC shows increased connectivity.

**Figure 5:**
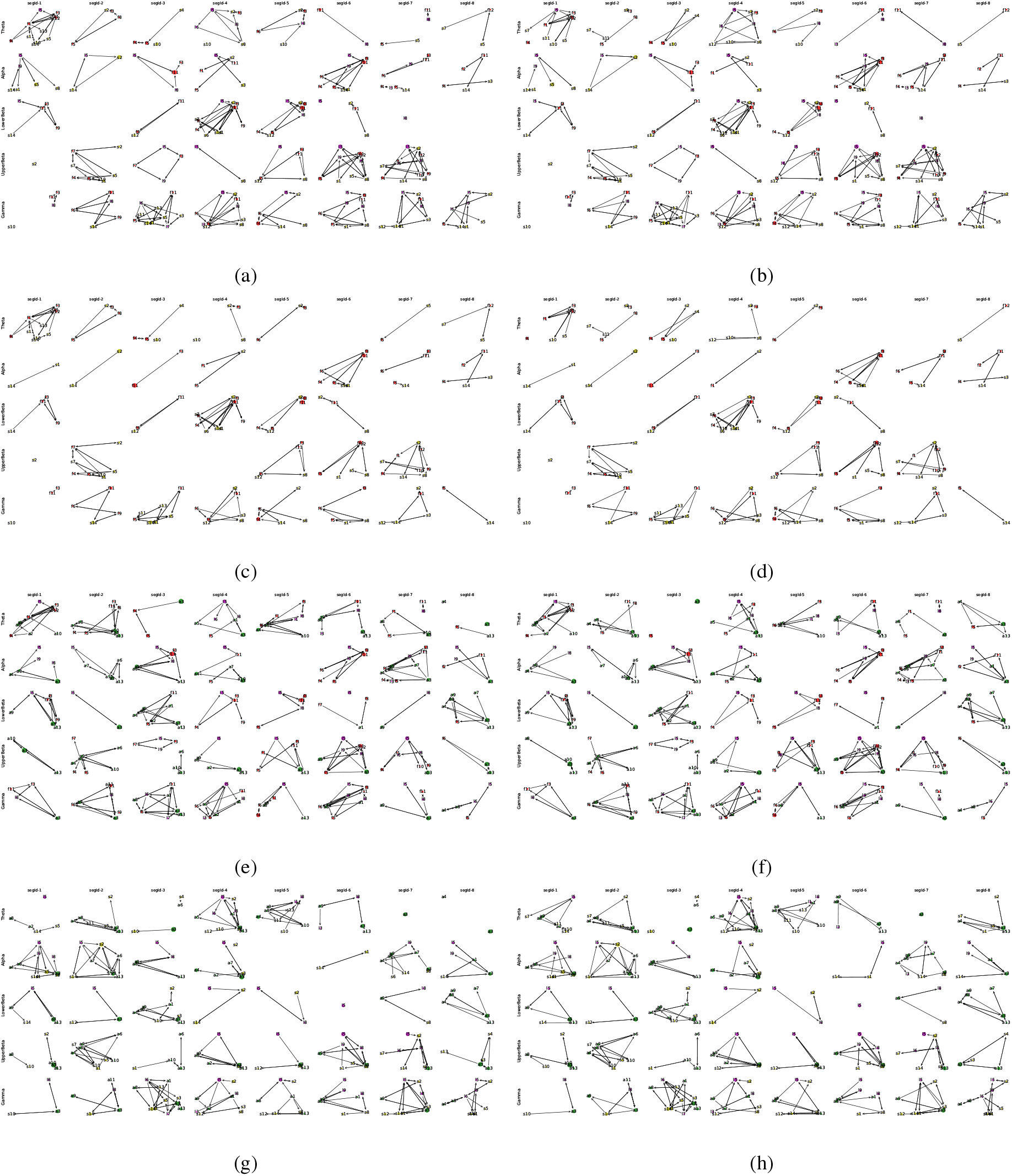
All the graphs are created for the representative (median) connections across the high-familiarity (first column) and less-familiarity (second column) groups. (a-b) For language, core Affect and exteroceptive sensory related regions. (c-d) For core Affect and exteroceptive sensory related regions. (e-f) For language, core Association and core Affect networks. (g-h) For language, core Association and exteroceptive sensory regions. For encoding of regions see supplementary table-1

**Figure 6:**
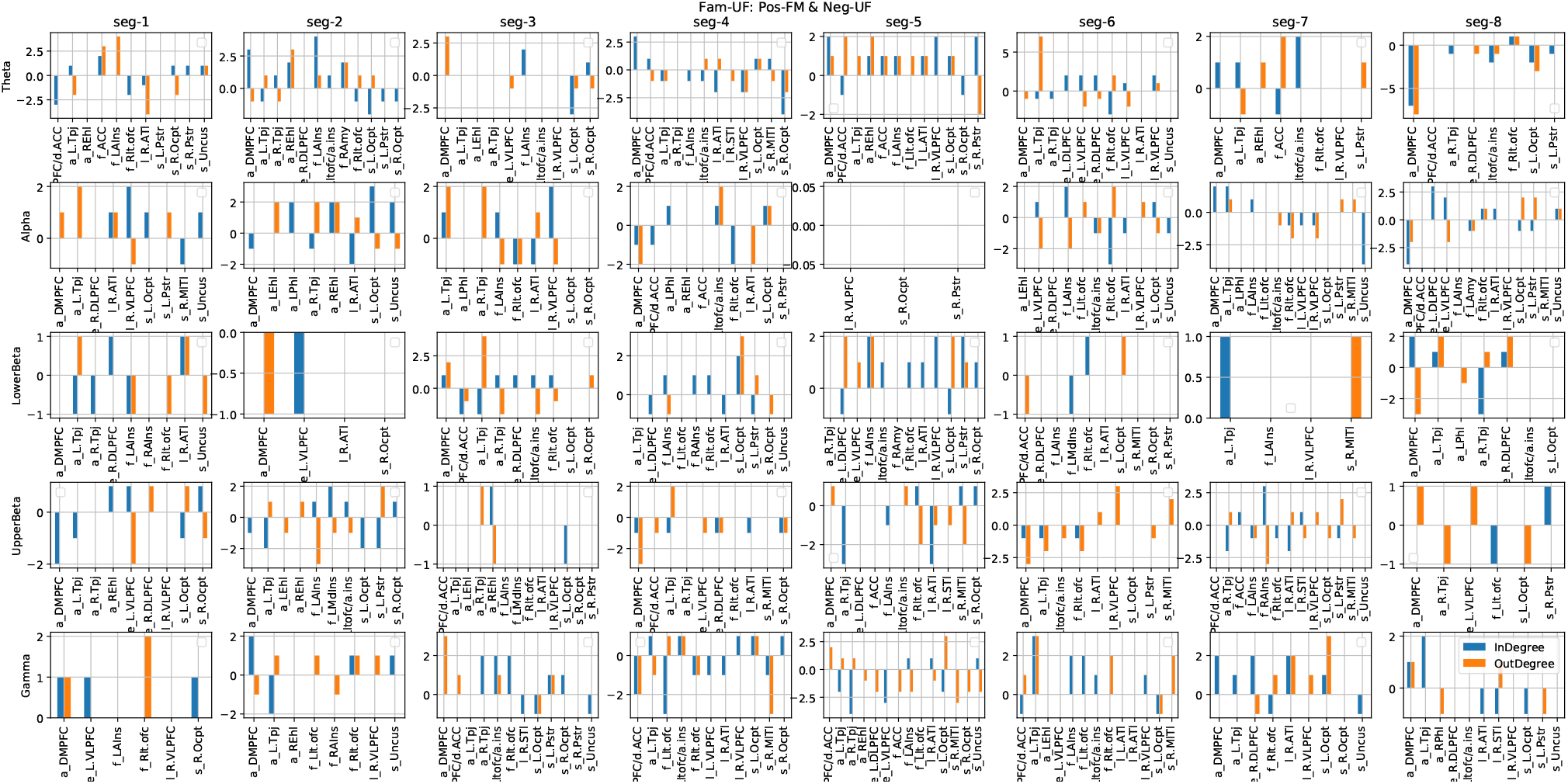
The difference between in-degree and out-degree for less-familiar and high-familiar stimuli is calculated. In the y-axis, positive values indicates when high-familiarity is greater and vice-versa for the negative values. x-axis represents name of the regions. The format for the region names is *< networkname >* _ *< laterality >< region abbr. >*. The difference is calculated for all the frequency bands (in row) and for all the segments (in column).

OFC receives highly processed sensory information, including information about bodily states such as hunger and thirst and inputs from areas that process high-level emotional and social information and connects with the insula, amygdala, and other parts of the prefrontal cortex [47]. These connections enable OFC to encode associations between sensory stimuli in the external world and internal states related to emotionally relevant events. For instance, activity in OFC is reported when participants are tracking dynamically changing emotional conditions based on socio-emotional information [16]. In addition, the role of OFC is also reported in the valence tracking [16]. Furthermore, it is shown that during reward uncertainty, the activity in the OFC plays a vital role in encoding reward-biased and risk-modulated subjective value dynamically [27] hence, facilitates the flexible adaptation to the circumstances [10, 1]. These evidence suggest that increased connectivity of OFC with other high-level regions facilitates the adaptation and valence tracking in the relatively less familiar environment. On the other hand, our results observe comparatively more connectivity of dmPFC for less-familiar contexts. Activity in the dmPFC is mostly reported in resolving valence & emotional conflicts [28, 21, 13, 39] and in dynamic adaptation to stimulus-response values [36, 15].

Whereas, increased insular connectivity mostly in lower beta and gamma bands is observed for high-context familiarity (fig-6 & 4). It is reported by [7] that neural activity in the beta and gamma bands is distinctly associated with topdown and bottom-up streams. AIC is involved in integrating bottom-up interoceptive signals with top-down predictions. The descending predictions to visceral systems provide a point of reference for autonomic reflexes[19]. In addition, the frontotemporal connections of the insular network play a significant role in social cognition, emotion and interoception. For instance, a meta-analysis study on the healthy population and lesion studies [2] concluded that the insula and its extended network with frontotemporal regions are important for a triangle of social cognition, emotion and interoception. Another review study on Insula [12] reported the involvement of AIC in emotion awareness and feelings. In our power-correlation analysis of ECG with different brain networks(fig-2a), we found that cardiac activity is correlated with the core-affect network-related regions during seg-3 to seg-7, which is resonating with previous studies that the insular cortex with cortical regions including ACC is involved in regulating cardiac signals[40].

Furthermore, we observed connectivity in ATL regions for both high-familiar and less-familiar context stimuli. Anterior temporal lobes are generally involved in social cognition, theory of mind processing, social conceptual knowledge, and memory systems that are linked to reward and valence[22]. ATL is reported as the multimodal integration hub, which integrates information from different modalities, including internal body physiology, exteroceptive sensations, motivation and hedonic input, and social cognition to represent the context-sensitive conceptual knowledge and semantic meaning. Conceptual representations are distilled within the system, which encodes knowledge of concepts by learning the higher-order relationships among various sensory, motor, linguistic, social, and affective sources of information widely distributed in the cortex. This higher-order learning is a lifelong process cultured through verbal and non-verbal embodied experiences [33, 32, 43] in order to conceptualize the abstract knowledge across contexts and items with their context-based controlled access [46, 42, 44,43].

## 5. Conclusion

In summary, in this work, we have tried to address how the varying degrees of certainty about the context can influence affective feelings, brain dynamics, and cardiac-brain interaction. To address the question, we adapted the naturalistic emotion paradigm and tracked the brain dynamics & cardiac-brain activity during pre-awareness to consciously feeling state. With this work, we also provided empirical support to the computational model of valence [23] and active interoceptive inference (AII) model [3]. The experiment, analysis, and results presented here contribute significantly to the understanding of brain dynamics and cardiac-brain interaction concerning the degree of emotional context un/familiarity.

## References

[1] Abela, A.R., Chudasama, Y., 2013. Dissociable contributions of the ventral hippocampus and orbitofrontal cortex to decision-making with a delayed or uncertain outcome. European Journal of Neuroscience 37, 640–647.

[2] Adolfi, F., Couto, B., Richter, F., Decety, J., Lopez, J., Sigman, M., Manes, F., Ibanez, A., 2017. Convergence of interoception, emotion, and social cognition: a twofold fmri meta-analysis and lesion approach. Cortex 88, 124–142.

[3] Allen, M., Levy, A., Parr, T., Friston, K.J., 2019. In the bodys eye: the computational anatomy of interoceptive inference. BioRxiv, 603928.

[4] Anderson, E., Siegel, E.H., Bliss-Moreau, E., Barrett, L.F., 2011. The visual impact of gossip. Science 332, 1446–1448.

[5] Anderson, E.C., Carleton, R.N., Diefenbach, M., Han, P.K., 2019. The relationship between uncertainty and affect. Frontiers in psychology 10, 2504.

[6] Azevedo, R.T., Badoud, D., Tsakiris, M., 2018. Afferent cardiac signals modulate attentional engagement to low spatial frequency fearful faces. Cortex 104, 232–240.

[7] Bastos, A.M., Vezoli, J., Bosman, C.A., Schoffelen, J.M., Oostenveld, R., Dowdall, J.R., De Weerd, P., Kennedy, H., Fries, P., 2015. Visual areas exert feedforward and feedback influences through distinct frequency channels. Neuron 85, 390–401.

[8] Baumeister, R.F., Bratslavsky, E., Finkenauer, C., Vohs, K.D., 2001. Bad is stronger than good. Review of general psychology 5, 323–370.

[9] Carleton, R.N., 2016. Into the unknown: A review and synthesis of contemporary models involving uncertainty. Journal of anxiety disorders 39, 30–43.

[10] Chau, B.K., Sallet, J., Papageorgiou, G.K., Noonan, M.P., Bell, A.H., Walton, M.E., Rushworth, M.F., 2015. Contrasting roles for orbitofrontal cortex and amygdala in credit assignment and learning in macaques. Neuron 87, 1106–1118.

[11] Corcoran, A.W., Macefield, V.G., Hohwy, J., 2021. Be still my heart: Cardiac regulation as a mode of uncertainty reduction. Psychonomic Bulletin & Review, 1–13.

[12] Craig, A.D., Craig, A., 2009. How do you feel–now? the anterior insula and human awareness. Nature reviews neuroscience 10.

[13] De Wit, S., Kosaki, Y., Balleine, B.W., Dickinson, A., 2006. Dorsomedial prefrontal cortex resolves response conflict in rats. Journal of Neuroscience 26, 5224–5229.

[14] Dieterich, R., Endrass, T., Kathmann, N., 2016. Uncertainty is associated with increased selective attention and sustained stimulus processing. Cognitive, Affective, & Behavioral Neuroscience 16, 447–456.

[15] Gonzalez-Escamilla, G., Chirumamilla, V.C., Meyer, B., Bonertz, T., von Grotthus, S., Vogt, J., Stroh, A., Horstmann, J.P., Tüscher, O., Kalisch, R., et al., 2018. Excitability regulation in the dorsomedial prefrontal cortex during sustained instructed fear responses: a tmseeg study. Scientific reports 8, 1–12.

[16] Goodkind, M.S., Sollberger, M., Gyurak, A., Rosen, H.J., Rankin, K.P., Miller, B., Levenson, R., 2012. Tracking emotional valence: the role of the orbitofrontal cortex. Human brain mapping 33, 753–762.

[17] Gottlieb, J., Hayhoe, M., Hikosaka, O., Rangel, A., 2014. Attention, reward, and information seeking. Journal of Neuroscience 34, 15497–15504.

[18] Gray, J., McNaughton, N., 2000. The neuropsychology of anxiety: An enquiry into the functions of the septo-hippocampal system. 2nd edn oxford university press.

[19] Gu, X., Hof, P.R., Friston, K.J., Fan, J., 2013. Anterior insular cortex and emotional awareness. Journal of Comparative Neurology 521, 3371–3388.

[20] Guo, C.C., Nguyen, V.T., Hyett, M.P., Parker, G.B., Breakspear, M.J., 2015. Out-of-sync: disrupted neural activity in emotional circuitry during film viewing in melancholic depression. Scientific reports 5, 1–12.

[21] Haddon, J.E., Killcross, A.S., 2005. Medial prefrontal cortex lesions abolish contextual control of competing responses. Journal of the experimental analysis of behavior 84, 485–504.

[22] Hertrich, I., Dietrich, S., Ackermann, H., 2020. The margins of the language network in the brain. Frontiers in Communication 5, 93.

[23] Hesp, C., Smith, R., Parr, T., Allen, M., Friston, K.J., Ramstead, M.J., 2021. Deeply felt affect: The emergence of valence in deep active inference. Neural Computation 33, 1–49.

[24] Hirsh, J.B., Mar, R.A., Peterson, J.B., 2012. Psychological entropy: a framework for understanding uncertainty-related anxiety. Psychological review 119, 304.

[25] Hofmann, S.M., Klotzsche, F., Mariola, A., Nikulin, V.V., Villringer, A., Gaebler, M., 2020. Decoding subjective emotional arousal from eeg during an immersive virtual reality experience. bioRxiv.

[26] Hulme, O.J., Morville, T., Gutkin, B., 2019. Neurocomputational theories of homeostatic control. Physics of life reviews 31, 214–232.

[27] Jo, S., Jung, M.W., 2016. Differential coding of uncertain reward in rat insular and orbitofrontal cortex. Scientific reports 6, 1–13.

[28] Kotz, S.A., Dengler, R., Wittfoth, M., 2015. Valence-specific conflict moderation in the dorso-medial pfc and the caudate head in emotional speech. Social cognitive and affective neuroscience 10, 165–171.

[29] Lindquist, K.A., Wager, T.D., Kober, H., Bliss-Moreau, E., Barrett, L.F., 2012. The brain basis of emotion: a meta-analytic review. The Behavioral and brain sciences 35, 121.

[30] Ludwig, R.J., Welch, M.G., 2020. How babies learn: the autonomic socioemotional reflex. Early Human Development, 105183.

[31] Lynn, S.K., Barrett, L.F., 2014. utilizing signal detection theory. Psychological science 25, 1663–1673.

[32] Martin, A., 2016. Grapesgrounding representations in action, perception, and emotion systems: How object properties and categories are represented in the human brain. Psychonomic bulletin & review 23, 979–990.

[33] Meteyard, L., Cuadrado, S.R., Bahrami, B., Vigliocco, G., 2012. Coming of age: A review of embodiment and the neuroscience of semantics. Cortex 48, 788–804.

[34] Mishra, S., Asif, M., Tiwary, U.S., 2021a. Dataset on emotions using naturalistic stimuli (dens). bioRxiv URL: https://www.biorxiv.org/content/early/2021/08/05/2021.08.04.455041, doi:10.1101/2021.08.04.455041, arXiv:https://www.biorxiv.org/content/early/2021/08/05/2021.08.04.455041.full.pd

[35] Mishra, S., Asif, M., Tiwary, U.S., 2021b. A study of spatio-temporal dynamics of emotion processing usingdens dataset. bioRxiv URL: https://www.biorxiv.org/content/early/2021/08/05/2021.08.05.455187, doi:10.1101/2021.08.05.455187, arXiv:https://www.biorxiv.org/content/early/2021/08/05/2021.08.05.455187.full.pd

[36] Mitchell, D.G., Luo, Q., Avny, S.B., Kasprzycki, T., Gupta, K., Chen, G., Finger, E.C., Blair, R.J.R., 2009. Adapting to dynamic stimulus-response values: differential contributions of inferior frontal, dorsomedial, and dorsolateral regions of prefrontal cortex to decision making. Journal of Neuroscience 29, 10827–10834.

[37] Moors, A., Ellsworth, P.C., Scherer, K.R., Frijda, N.H., 2013. Appraisal theories of emotion: State of the art and future development. Emotion Review 5, 119–124.

[38] Nguyen, V.T., Sonkusare, S., Stadler, J., Hu, X., Breakspear, M., Guo, C.C., 2017. Distinct cerebellar contributions to cognitive-perceptual dynamics during natural viewing. Cerebral Cortex 27, 5652–5662.

[39] Oehrn, C.R., Hanslmayr, S., Fell, J., Deuker, L., Kremers, N.A., Do Lam, A.T., Elger, C.E., Axmacher, N., 2014. Neural communication patterns underlying conflict detection, resolution, and adaptation. Journal of Neuroscience 34, 10438–10452.

[40] Oppenheimer, S., Cechetto, D., 2011. The insular cortex and the regulation of cardiac function. Comprehensive Physiology 6, 1081–1133.

[41] Pascual-Marqui, R.D., et al., 2002. Standardized low-resolution brain electromagnetic tomography (sloreta): technical details. Methods Find Exp Clin Pharmacol 24, 5–12.

[42] Patterson, K., Nestor, P.J., Rogers, T.T., 2007. Where do you know what you know? the representation of semantic knowledge in the human brain. Nature reviews neuroscience 8, 976–987.

[43] Ralph, M.A.L., Jefferies, E., Patterson, K., Rogers, T.T., 2017. The neural and computational bases of semantic cognition. Nature Reviews Neuroscience 18, 42–55.

[44] Ralph, M.A.L., Sage, K., Jones, R.W., Mayberry, E.J., 2010. Coherent concepts are computed in the anterior temporal lobes. Proceedings of the National Academy of Sciences 107, 2717–2722.

[45] Rao, V.R., Sellers, K.K., Wallace, D.L., Lee, M.B., Bijanzadeh, M., Sani, O.G., Yang, Y., Shanechi, M.M., Dawes, H.E., Chang, E.F., 2018. Direct electrical stimulation of lateral orbitofrontal cortex acutely improves mood in individuals with symptoms of depression. Current Biology 28, 3893–3902.

[46] Rogers, T.T., Lambon Ralph, M.A., Garrard, P., Bozeat, S., McClelland, J.L., Hodges, J.R., Patterson, K., 2004. Structure and deterioration of semantic memory: a neuropsychological and computational investigation. Psychological review 111, 205.

[47] Rudebeck, P.H., Rich, E.L., 2018. Orbitofrontal cortex. Current Biology 28, R1083–R1088.

[48] Ruiz-Padial, E., Vila, J., Thayer, J.F., 2011. The effect of conscious and non-conscious presentation of biologically relevant emotion pictures on emotion modulated startle and phasic heart rate. International Journal of Psychophysiology 79, 341–346.

[49] Salvi, C., Beeman, M., Bikson, M., McKinley, R., Grafman, J., 2020. Tdcs to the right anterior temporal lobe facilitates insight problemsolving. Scientific reports 10, 1–10.

[50] Schubring, D., Schupp, H.T., 2019. Affective picture processing: Alpha-and lower beta-band desynchronization reflects emotional arousal. Psychophysiology 56, e13386.

[51] Seth, A.K., Friston, K.J., 2016. Active interoceptive inference and the emotional brain. Philosophical Transactions of the Royal Society B: Biological Sciences 371, 20160007.

[52] Silvani, A., Calandra-Buonaura, G., Dampney, R.A., Cortelli, P., 2016. Brain–heart interactions: physiology and clinical implications. Philosophical Transactions of the Royal Society A: Mathematical, Physical and Engineering Sciences 374, 20150181.

[53] Tan, A.Y., Chen, Y., Scholl, B., Seidemann, E., Priebe, N.J., 2014. Sensory stimulation shifts visual cortex from synchronous to asynchronous states. Nature 509, 226–229.

[54] Thayer, J.F., Hansen, A.L., Saus-Rose, E., Johnsen, B.H., 2009. Heart rate variability, prefrontal neural function, and cognitive performance: the neurovisceral integration perspective on self-regulation, adaptation, and health. Annals of Behavioral Medicine 37, 141–153.

[55] Viinikainen, M., Glerean, E., Jääskeläinen, I.P., Kettunen, J., Sams, M., Nummenmaa, L., 2012. Nonlinear neural representation of emotional feelings elicited by dynamic naturalistic stimulation. Open Journal of Neuroscience 2, 1–17.

[56] Vuilleumier, P., Armony, J.L., Driver, J., Dolan, R.J., 2003. Distinct spatial frequency sensitivities for processing faces and emotional expressions. Nature neuroscience 6, 624–631.

[57] Waschke, L., Tune, S., Obleser, J., 2019. Local cortical desynchronization and pupil-linked arousal differentially shape brain states for optimal sensory performance. Elife 8, e51501.

[58] Wass, S., De Barbaro, K., Clackson, K., 2016. Learning and the autonomic nervous system: understanding interactions between stress, concentration and learning during early childhood, in: Front. Neurosci. doi: 10.3389/conf.fnins.

[59] Zerlaut, Y., Destexhe, A., 2017. Enhanced responsiveness and lowlevel awareness in stochastic network states. Neuron 94, 1002–1009.

