## supplementary fig-1, supplementary fig-2, supplementary fig-3, supplementary fig-4, supplementary table-1 for "The degree of context un/familiarity impacts the emotional feeling and preaware cardiac-brain activity: a study with emotionally salient naturalistic paradigm using DENS Dataset"

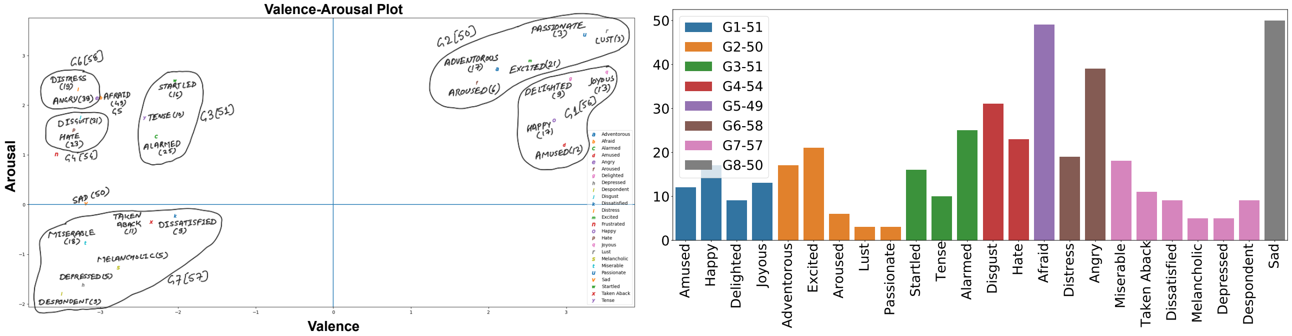


(a) Valence Arousal Grouping

(b) Histogram of Emotion Grouping

**Figure 1: SI:Groupings** We have grouped the emotions based on two criteria. First, the distance-based proximity on V-A space, and second, approximately equal samples across emotion groups. (b) Histogram depicting the count of individual emotions and their assignment to different emotion groups.

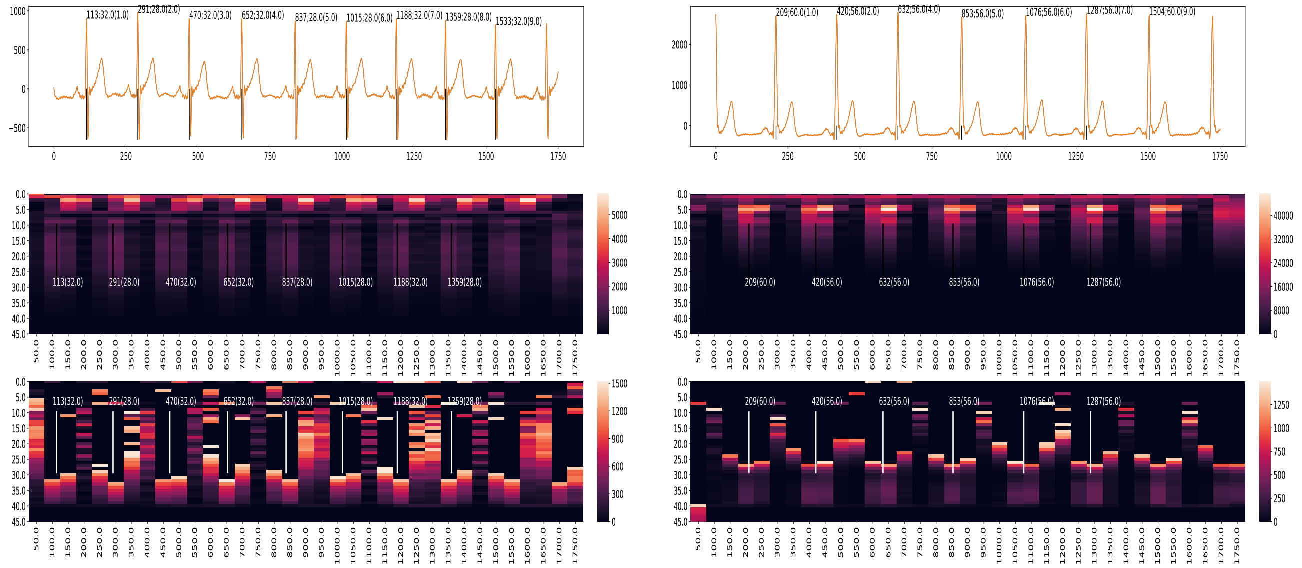

(a)

(b)

**Figure 2: SI:ECG** (a-b) ECG with spectrogram calculated using stft. First diagram in (a-b) is ECG graph, second diagram is spectrogram calculated using stft, and third diagram is emphasizing low magnitude frequencies ($mag > 450$). The annotation in ECG graph is time of Rpeak:duration of Rpeak(segment number). For low R-width, the magnitude of higher frequencies is relatively more than the higher frequency magnitude for high R-width (as shown in the graph relatively more magnitude for 15Hz to 30 Hz for low R-width).

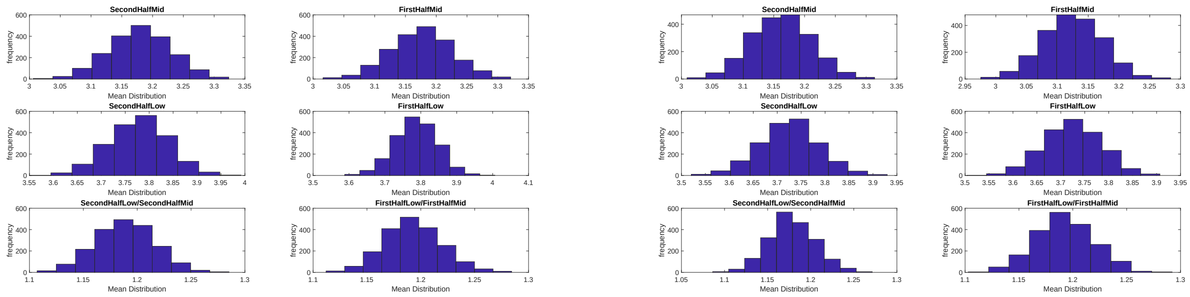


(a)

(b)

**Figure 3: SI:Mean distribution** Mean distribution plots for (a) High-Familiar (b) Less-Familiar.

| Node | RegName |
|---|---|
| a1 | DMPFC/d.ACC |
| a10 | REhl |
| a11 | REhl |
| a12 | RPhl |
| a13 | R.Tpj |
| a2 | DMPFC/d.ACC |
| a3 | DMPFC |
| a4 | DMPFC |
| a5 | DMPFC |
| a6 | LEhl |
| a7 | LPhl |
| a8 | L.Tpj |
| a9 | L.Tpj |
| e1 | L.DLPFC |
| e2 | L.VLPFC |
| e3 | R.DLPFC |
| f1 | ACC |
| f10 | RAIns |
| f11 | Rlt.ofc |
| f12 | Rlt.ofc |
| f2 | LAmy |
| f3 | LAIns |
| f4 | LAIns |
| f5 | Lltofc/a.ins |
| f6 | Llt.ofc |
| f7 | LMdIns |
| f8 | RAmy |
| f9 | RAIns |
| l1 | L.ATl |
| l3 | L.VLPFC |
| l4 | R.ATl |
| l5 | R.ATl |
| l6 | R.STl |
| l7 | R.STl |
| l8 | R.VLPFC |
| l9 | R.VLPFC |
| s1 | L.Ocpt |
| s10 | R.Ocpt |
| s11 | R.Ocpt |
| s12 | R.Ocpt |
| s13 | R.Pstr |
| s14 | Uncus |
| s2 | L.Ocpt |
| s3 | L.Ocpt |
| s4 | L.Ocpt |
| s5 | L.Pstr |
| s6 | L.Pstr |
| s7 | L.Pstr |
| s8 | R.MITl |

**Table 1**
**SI:Abbreviations** Abbreviations of network nodes. In the first column, the first letter in all row is: f-coreAffect, a-coreAssociation, l-language, s-exteroceptive network, e-executiveControl.

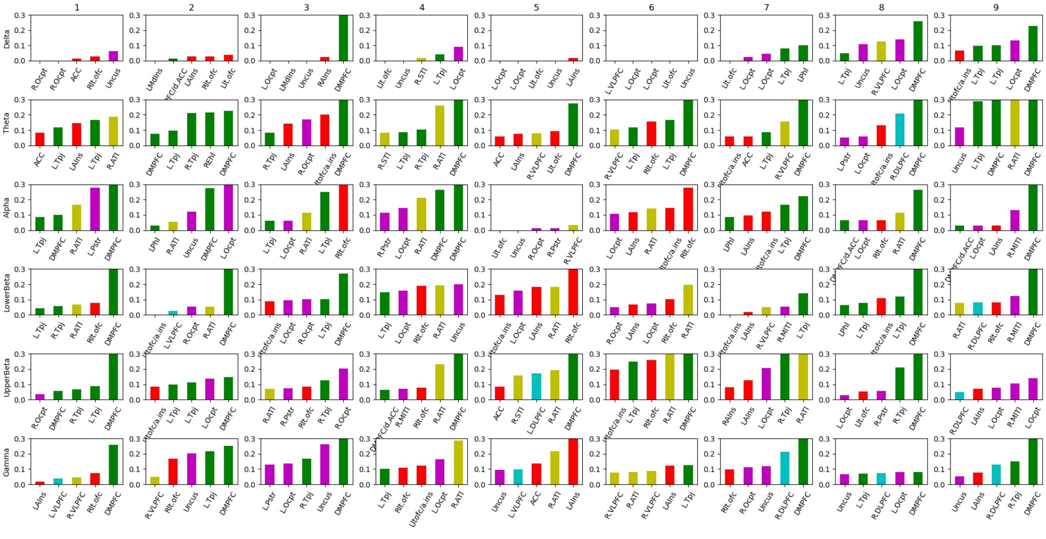

(a)

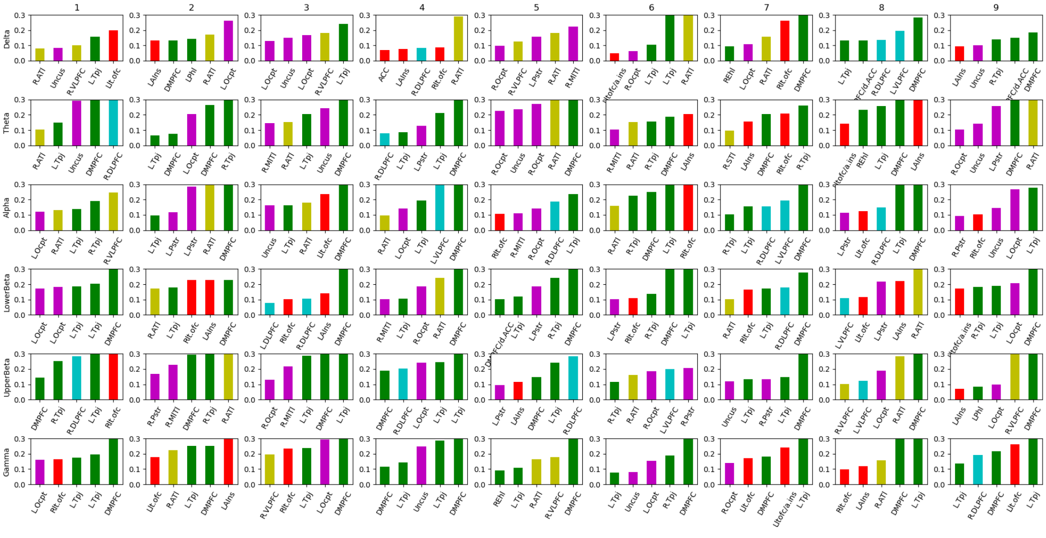

(b)

**Figure 4: SI:Hubs** (a) For emotion groups, (b) for eyes-open baseline state. seg wise hub dynamics while all the emotion groups are combined. Top-five hubs are displayed. These hubs are calculated using Betweenness centrality and responsible for integrating information. Green: core Association network; Red: core Affect network; Cyan: exectiveControl network; Magenta: extroception network; Yellow: Language network.
